# Varying trypsin inhibitor activity in differently processed soybean expellers linearly reduces prececal amino acid digestibility in broilers

**DOI:** 10.1101/2020.11.24.395988

**Authors:** S. Kuenz, S. Thurner, D. Hoffmann, K. Kraft, M. Wiltafsky-Martin, K. Damme, W. Windisch, D. Brugger

## Abstract

The present study investigated the effect of varying trypsin inhibitor activity (TIA) in differently processed soybean expellers on apparent prececal amino acid (AA) digestibility in male broiler chickens. Two different raw soybean batches were treated using four different processing techniques (thermal, hydrothermal, pressure, kilning) at varying intensities. In this way, 45 expeller extracted soybean meal (ESBM) variants were created. The processed soybean variants were then merged into a basal diet (160 g/kg crude protein (CP)) at two inclusion levels (15%, 30%) resulting in 91 different diets (1 basal diet plus 90 experimental diets) with TIA ranging from 0.4 mg/g to 8.5 mg/g. All diets contained 0.5% of titanium dioxide (TiO_2_). During four experimental runs, a total of 5,460 1-day old male broiler chickens (Ross 308) were fed a commercial starter diet (CP 215 g/kg, 14 g/kg Lysine, 12.5 MJ ME/kg) ad libitum from day 1 to day 14. Subsequently, birds were allocated to a total of 546 pens with 10 birds per pen and were fed the 91 experimental diets *ad libitum*. At day 22, birds were sacrificed and digesta of the terminal ileum was collected for determination of AA digestibility. TIA depressed the prececal digestibility of every single AA significantly in a straight linear fashion (p < 0.001). cystine and methionine expressed the strongest suppression by TIA with cystine showing the lowest apparent prececal digestibility measured (4.94% at 23.6 mg/g TIA in raw ESBM). Correspondingly, live weight (LW) (p < 0.001) and total weight gain (TWG) (p < 0.001) declined in a linear manner with increasing TIA in feed. The present data demonstrate that TIA severely depresses digestibility of essential and non-essential AA and thus growth performance in a straight linear fashion. On the one hand, this questions the usefulness of defined upper limits of TIA in soy products whereas on the other hand, TIA must be considered when testing raw components for their feed protein value in vivo.

## 1. Introduction

Soybeans are the most important protein source in livestock feeding. Compared to other plants, soybeans have high contents of oil and protein and a superior amino acid pattern (Clarke and Wiseman, 2005). The nutritional value of most plant materials is limited by the presence of numerous naturally occurring compounds, which interfere with nutrient digestion and absorption (Clarke and Wiseman, 2000). The most important anti-nutritional substances occurring in soybeans are trypsin inhibitors (TI), which can be divided into two classes: The Kunitz trypsin inhibitors (KTI) and the Bowman-Birk trypsin inhibitors (BBI). The KTI consist of 181 amino acid residues, have a relative molecular weight of 20,100 Da (Koide and Ikenaka, 1973). The reactive site of this inhibitor class is located at residues Arg 63 and Ile 64 and binds primarily trypsin. The KTI have a low content of cysteine and only two disulfide bonds. The second class, BBI are rich in cysteine and have seven disulfide bridges, which is the reason for a very dense three-dimensional structure (Odani and Ikenaka, 1973). In comparison to the KTI, the BBI are relatively low in molecular weight (approx. 6,000 Da to 10,000 Da) and have two independent and symmetric binding sites for trypsin and chymotrypsin. The trypsin-reactive site is located on Lys 16 and Ser 17 and the chymotrypsin-reactive site is positioned on Leu 43 and Ser 44 (Odani and Ikenaka, 1973). Both inhibitor classes form stable enzyme-inhibitor complexes on a molar 1:1 ratio (Clarke and Wiseman, 2000).

Earlier studies have been published showing the negative effect of trypsin inhibitor activity (TIA) on growth performance of rats (Grant et al., 1995; Gu et al., 2010), chickens (Clarke and Wiseman, 2007, 2005; Heger et al., 2016), turkeys (Mian and Garlich, 1995) and pigs (Batterham et al., 1993; Herkelman et al., 1992; Zollitsch et al., 1993). Furthermore, TI causes pancreas hypertrophy and hyperplasia in rats (Abbey et al., 1979; Grant et al., 1995) and chickens (Gertler et al., 1967; Hoffmann et al., 2019; Pacheco et al., 2014; Perilla et al., 1997). In response to the inhibition of digestive processes, the pancreas increases its size and number of acinar cells in order to elevate the secretion of digestive enzymes (Nitsan and Liener, 1976). In this context, it seems like the underlying endocrine signal that facilitates these adaptions is gut-derived cholecystokinin (CCK), which responds to a lower influx of free amino acids (AA) into the enterocytes of the small intestine (Miura et al., 1997). Due to the increased activity of the pancreas, Lyman and Lepkovski (1957) suggested that the overall depression in growth performance is mainly due to the high endogenous loss of AA and enzymes secreted from the pancreas, since the digestive depression itself could be quite efficiently compensated by the higher pancreatic enzyme secretion.

To avoid depression in performance, the heat-labile anti-nutritional factors in soybean products have to be sufficiently deactivated. Batterham et al. (1993) recommended to reduce TIA in soybean products for growing pigs to a level of 4.7 mg/g. Similar results were shown in the studies of Clarke and Wiseman (2007, 2005) for poultry. They concluded that TIA in full-fat soybeans should not exceed 4.0 mg/g. On the other hand, it is important to avoid protein denaturation induced heat damage. Pacheco et al. (2014) and Araba and Dale (1990) observed a decline in performance with decreasing protein solubility in potassium hydroxide (KOH) below 74% and 70%, respectively, in response to the heat-associated peptide denaturation. Another indicator for heat damage is the concentration of reactive lysine (Fontaine et al., 2007). The ϵ-amino group of lysine binds irreversibly with reducing sugars during heat treatment. The balance of adequate denaturation of TI and heat damage is controversial and discussed in literature: Heger et al. (2016) concluded in their study that growth performance is not impaired even above TIA of 4.0 mg/g, whereas Hoffmann et al. (2019) observed a linear improvement in feed efficiency when gradually decreasing TIA below 1.0 mg/g without impairing growth performance through excessive heat-damage to the protein fraction.

The parameter of choice for the determination of the feed protein value is the AA flow at the terminal ileum (Ravindran et al., 1999). This consists of the undigested and unabsorbed feed-borne AA as well as endogenously secreted AA. The pool of endogenously secreted AA is further subcategorized into the basal and specific endogenous losses. While the basal losses are considered to be predominantly affected by the total dry matter intake, the specific AA secretion appears to be affected by the quantity and characteristics of the protein under study (Angkanaporn et al., 1997; Dänicke et al., 2000; Souffrant, 2001). Hence, when determining feed protein quality as a function of AA flow at the terminal ileum, the specific endogenous AA losses should be considered. However, attempts to quantify endogenous protein secretion yield highly variable results (Donkoh and Moughan, 1999). Therefore, Rodehutscord et al. (2004) proposed an approach by which the digestibility of the dietary protein until the end of the ileum (so called “prececal digestibility”) is estimated through linear regression, as the slope of increasing apparently digested AAs until the terminal ileum with gradually increasing dietary AA intake via the feed protein under study. This method is supposed to exclude effects of varying endogenous protein losses, since the slope represents the prececal AA digestibility as affected by the protein source under study.

This study aimed to investigate the effects of differentially treated expeller extracted soybean meal (ESBM) and associated finely graded differences in dietary TIA in the feeding of broilers. In particular, the prececal amino acid digestibility according to Rodehutscord et al. (2004) as well as the zootechnical performance of birds were investigated.

## 2. Materials and Methods

### 2.1 Soybean processing and diet composition

Raw material for soy processing consisted of two homogenous batches of soybeans. In Batch 1 (breed: Sultana, native TIA: 37.3 mg/g) were conventionally produced soybeans, harvested in Germany, Batch 2 (breed: Merlin, native TIA: 40.5 mg/g) consisted of organically produced soybeans from Romania. These two batches were treated equally using four different processing techniques (thermal, hydrothermal, pressure and kilning) at varying processing intensities.

For the thermal treatment, soybeans were moistened and toasted for 40 seconds at either 115°C or 120°C, respectively. The hydrothermal method comprised the usage of steam at an average temperature of 103°C for about 40 minutes. The third method included hydrothermal treatment in combination with expander extrusion at intensities varying from 110°C to 130°C for at least one second to a maximum of five seconds. Using the kilning method, anti-nutritional factors were heat inactivated by hot recirculating air at 130°C up to 190°C with varying duration from 20 to 40 minutes. After cooling, all differently treated soybeans were mechanically de-oiled. In this way 45 differently ESBM were created. Table 1 presents the different treatments in detail, which have been already described earlier by Hoffmann et al. (2017). The intention with this processing scheme was to create a wide range of TIA and parameters associated with potential heat-damage to the protein fraction (KOH solubility, reactive lysine) to investigate the anti-nutritional effect of TIA and its potential interaction with protein damage on the AA utilization within the small intestine. The processing in general was not the objective of the present study, which experimental design would be not appropriate to investigate effects arising from different processing plants. The TIA in ESBM variants ranged from 0.3 mg/g to 23.6 mg/g, KOH soluble crude protein (CP) varied from 64.4% to 97.7% of total CP and reactive lysine from 14.7% to 25.0% of DM. TIA and heat-damage parameters, as well as AA concentrations are presented in Table 2. The processed ESBM were then merged into the basal diet (Table 3) at either 15% (CP level 1) or 30% (CP level 2) at the expense of maize starch according to Rodehutscord et al. (2004). Thereby, 91 experimental diets were mixed and subsequently pelletized to stabilize particle size distribution. All diets included 0.5% of titanium dioxide (TiO_2_) as an indigestible marker. The TIA in the final feed mixtures varied from 0.4 mg/g in the non-supplemented basal diet to 8.5 mg/g in the diet containing non-heated ESBM at the highest inclusion rate (30%).

**Table 1:** Soybean processing scheme for treatment technique and intensities adapted according to Hoffmann et al (2017)

| Thermal processing<br>[°C; min] | Hydrothermal processing | Pressure and thermal processing | Kilning and thermal processing |
| --- | --- | --- | --- |
|  | [ST: min, LT: min, Exp.: °C]<sup>1</sup> |  | [°C; min] |
| 115; 0.6 | ST: 0; LT: 00; Exp.: 0 | ST: 1; LT: 00; Exp.: 110 | 0; 0 |
| 120; 0.6 | ST: 1; LT: 03; Exp.: 0 | ST: 1; LT: 03; Exp.: 110 | 130; 40 |
|  | ST: 1; LT: 12; Exp.: 0 | ST: 1; LT: 03; Exp.: 130 | 140; 20 |
|  | ST: 1; LT: 48; Exp.: 0 | ST: 1; LT: 12; Exp.: 110 | 140; 30 |
|  |  | ST: 1; LT: 12; Exp.: 130 | 140; 40 |
|  |  | ST: 1; LT: 48; Exp.: 110 | 160; 30 |
|  |  | ST: 1; LT: 48; Exp.: 130 | 165; 20 |
|  |  |  | 165; 30 |
|  |  |  | 190; 20 |
|  |  |  | 190; 30 |

<sup>1</sup>ST: short time conditioning (90°C); LT: long time conditioning (100°C); Exp: expander

**Table 2:** Average chemical composition, crude protein quality and amino acid concentration, as well as range of processed expeller extracted soybean meals sorted according to treatment technique. All parameters are represented in percentage of dry matter, if not indicated otherwise

|  | Thermal processing |  | Hydrothermal processing |  | Pressure and thermal processing |  | Kilning and thermal processing |  |
| --- | --- | --- | --- | --- | --- | --- | --- | --- |
|  | Average | Range | Average | Range | Average | Range | Average | Range |
| *Chemical composition* |  |  |  |  |  |  |  |  |
| Dry matter | 96.9 ± 0.7 | 96.0- 97.8 | 87.2 ± 1.7 | 83.9- 90.4 | 87.1 ± 1.5 | 83.9- 89.0 | 95.6 ± 1.0 | 93.9- 96.9 |
| Crude protein | 49.9 ± 3.4 | 45.9- 53.0 | 47.3 ± 1.3 | 45.4- 49.3 | 47.2 ± 1.3 | 45.4- 49.3 | 45.7 ± 1.5 | 43.0- 47.7 |
| Crude ash | 6.4 ± 0.1 | 6.3- 6.5 | 6.4 ± 0.2 | 6.1- 6.7 | 6.4 ± 0.2 | 6.1- 6.7 | 6.2 ± 0.2 | 5.9- 6.7 |
| Crude fat | 10.0 ± 1.5 | 8.6- 11.9 | 10.8 ± 1.1 | 9.3- 13.1 | 10.4 ± 1.4 | 6.4- 13.7 | 12.1 ± 2.0 | 9.8- 15.4 |
| Crude fiber | 7.7 ± 1.6 | 6.3- 10.0 | 7.2 ± 0.6 | 6.3- 8.1 | 7.3 ± 0.6 | 6.3- 8.6 | 8.8 ± 1.5 | 7.3- 11.5 |
| *Crude protein quality* |  |  |  |  |  |  |  |  |
| TIA (mg/g DM) | 2.7 ± 0.9 | 1.9- 3.8 | 4.8 ± 7.1 | 0.3- 23.6 | 2.8 ± 4.0 | 0.3- 15.9 | 9.7 ± 4.2 | 5.5- 16.6 |
| KOH (% CP) | 81.6 ± 7.9 | 70.1- 88.0 | 83.5 ± 6.9 | 71.7- 97.6 | 82.1 ± 5.1 | 71.7- 91.8 | 79.3 ± 11.6 | 64.5- 95.4 |
| Total lysine (g/kg DM) | 25.9 ± 1.5 | 24.4- 28.0 | 23.6 ± 0.8 | 22.0- 25.0 | 23.4 ± 0.6 | 22.0- 25.0 | 23.4 ± 1.4 | 21.4- 25.7 |
| Reactive lysine (g/kg DM) | 21.0 ± 3.0 | 17.7- 25.0 | 18.5 ± 2.2 | 14.7- 22.0 | 17.9 ± 1.9 | 14.7- 22.0 | 17.4 ± 3.3 | 14.7- 22.0 |
| *Amino Acids* |  |  |  |  |  |  |  |  |
| Alanine | 2.04 ± 0.10 | 1.94- 2.16 | 1.74 ± 0.07 | 1.51- 1.81 | 1.75 ± 0.04 | 1.66- 1.81 | 1.87 ± 0.05 | 1.78- 1.92 |
| Arginine | 3.80 ± 0.36 | 3.43- 4.14 | 3.28 ± 0.20 | 2.59- 3.50 | 3.30 ± 0.12 | 3.09- 3.50 | 3.33 ± 0.14 | 3.12- 3.56 |
| Aspartic Acid | 5.53 ± 0.37 | 5.15- 5.90 | 4.69 ± 0.22 | 3.94- 4.95 | 4.73 ± 0.13 | 4.47- 4.95 | 4.96 ± 0.15 | 4.73- 5.16 |
| Cysteine | 0.68 ± 0.04 | 0.64- 0.72 | 0.60 ± 0.02 | 0.55- 0.63 | 0.60 ± 0.02 | 0.55- 0.63 | 0.64 ± 0.02 | 0.61- 0.68 |
| Glutamic Acid | 8.77 ± 0.72 | 8.14- 9.53 | 7.47 ± 0.40 | 6.15- 7.90 | 7.53 ± 0.26 | 7.03- 7.90 | 7.84 ± 0.22 | 7.50- 8.15 |
| Glycine | 2.01 ± 0.10 | 1.91- 2.12 | 1.73 ± 0.07 | 1.48- 1.82 | 1.74 ± 0.04 | 1.65- 1.82 | 1.84 ± 0.06 | 1.75- 1.91 |
| Histidine | 1.26 ± 0.08 | 1.18- 1.33 | 1.07 ± 0.05 | 0.91- 1.13 | 1.08 ± 0.03 | 1.02- 1.13 | 1.13 ± 0.04 | 1.08- 1.18 |
| Isoleucine | 2.18 ± 0.12 | 2.07- 2.29 | 1.85 ± 0.08 | 1.58- 1.95 | 1.86 ± 0.05 | 1.77- 1.95 | 1.97 ± 0.05 | 1.89- 2.03 |
| Leucine | 3.70 ± 0.22 | 3.49- 3.93 | 3.14 ± 0.14 | 2.66- 3.29 | 3.16 ± 0.08 | 2.99- 3.29 | 3.33 ± 0.09 | 3.19- 3.45 |
| Lysine | 2.93 ± 0.16 | 2.73- 3.10 | 2.56 ± 0.11 | 2.57- 2.71 | 2.58 ± 0.08 | 2.45- 2.71 | 2.62 ± 0.14 | 2.40- 2.84 |
| Methionine | 0.64 ± 0.03 | 0.61- 0.67 | 0.55 ± 0.02 | 0.49- 0.57 | 0.55 ± 0.01 | 0.52- 0.57 | 0.59 ± 0.02 | 0.57- 0.62 |
| Methionine + Cysteine | 1.33 ± 0.07 | 1.25- 1.39 | 1.15 ± 0.04 | 1.05- 1.19 | 1.15 ± 0.03 | 1.07- 1.19 | 1.24 ± 0.04 | 1.19- 1.29 |
| Phenylalanine | 2.46 ± 0.15 | 2.31- 2.61 | 2.09 ± 0.10 | 1.75- 2.21 | 2.10 ± 0.06 | 1.99- 2.21 | 2.21 ± 0.07 | 2.09- 2.30 |
| Proline | 2.42 ± 0.12 | 2.30- 2.53 | 2.06 ± 0.11 | 1.74- 2.22 | 2.07 ± 0.08 | 1.92- 2.22 | 2.18 ± 0.07 | 2.07- 2.28 |
| Serine | 2.45 ± 0.15 | 2.28- 2.61 | 2.08 ± 0.09 | 1.79- 2.18 | 2.10 ± 0.05 | 1.98- 2.18 | 2.21 ± 0.06 | 2.11- 2.29 |
| Threonine | 1.86 ± 0.09 | 1.77- 1.96 | 1.58 ± 0.06 | 1.38- 1.65 | 1.59 ± 0.03 | 1.52- 1.65 | 1.70 ± 0.05 | 1.63- 1.76 |
| Valine | 2.30 ± 0.15 | 2.17- 2.44 | 1.96 ± 0.09 | 1.66- 2.09 | 1.98 ± 0.05 | 1.88- 2.09 | 2.09 ± 0.06 | 1.99- 2.16 |

**Table 3:**
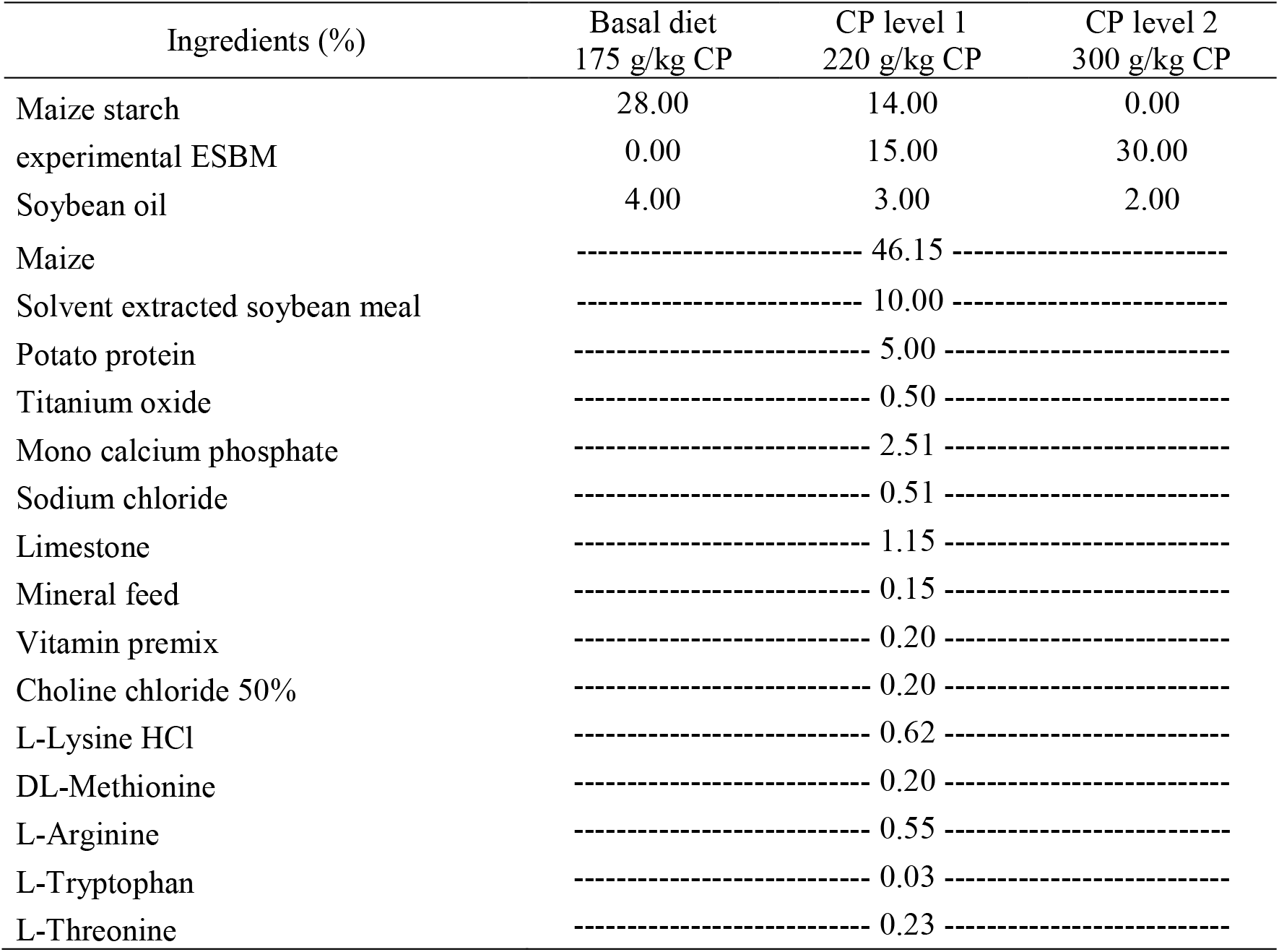
Feed composition of basal diet and experimental diets. The experimental diets contained two different levels of crude protein (CP level 1 = 220 g/kg CP and CP level 2 = 300 g/kg CP). This resulted in two varying inclusions of experimental expeller extracted soybean meal (ESBM) with an equivalent reduction of maize starch

### 2.2 Animals and experimental protocol

The study design was reviewed and approved by the responsible animal welfare officer of the Technical University of Munich and registered and approved by the legal authorities of the District Government of Lower Franconia, Federal State of Bavaria, Germany (registered case no. 55.2-DMS 2532-2-2164). The experiments took place at the Department for Education and Poultry Research, Bavarian State Research Center for Agriculture in Kitzingen (Germany).

The experiment was designed according to the model suggested by Rodehutscord et al. (2004). A total of 5,460 1-day-old male broiler chickens (Ross 308) were obtained from a local hatchery (Brüterei Süd, Regenstauf, Germany). Animals were reared with a commercial starter diet (CP 215 g/kg, 14 g/kg lysine, 12.5 MJ ME/kg) fed *ad libitum* from experimental day 1 to 14. On day 15, birds were weighed and randomly allocated to one of 546 pens (10 birds per pen, 1.6 m² per pen) equipped with feeder, nipple drinker and straw beddings. The 91 experimental diets were randomly distributed over pens, yielding an effective sample size of six replicates per feeding group. The diets were fed *ad libitum* until day 22 on which birds were weighed individually and euthanized by asphyxiation with carbon dioxide. The animals’ body cavities were immediately opened; the section between Meckel’s diverticulum and 2 cm anterior to the ileo-ceca-colonic-junction was isolated. The ileal content of two thirds of the terminal section was flushed with distilled water according to the method of Kluth et al. (2005). The digesta was pooled within each pen, frozen at −20°C and freeze-dried for later chemical analyses.

Throughout this study, birds had *ad libitum* access to drinking water (tap water) and the water consumption per pen was monitored continuously.

### 2.3 Chemical analyses

Raw soybean batches, ESBM and experimental diets were analyzed for TIA (DIN EN ISO 14902:2002-02) and crude nutrient values according to published procedures (VDLUFA, 2012). Furthermore, ESBM were analyzed for protein solubility in KOH (DIN EN ISO 14244:2014-02) and reactive lysine according to the homoarginine method applied by Pahm et al. (2008) to assess protein denaturation. AA content of ESBM, experimental diets and digesta were determined by ion-exchange chromatography referring to Llames and Fontaine (1994). According to this method, the oxidized cysteine- dimer cystine was measured. TiO_2_ in diets and digesta were analyzed according to the method of Brandt and Allam (1987). Analyzed crude nutrients and AA concentration of experimental diets are presented in Table 4.

**Table 4:** Mean values and standard deviation of analyzed nutrient concentration (% DM) and amino acid concentration (% DM) of experimental diets CP level 1 and CP level 2. Standard deviation of Basal diet refers to analytical replicate.

| Nutrient concentration (% DM) | Basal diet<br>160 g/kg CP | CP level 1<br>220 g/kg CP | CP level 2<br>300 g/kg CP |
| --- | --- | --- | --- |
| Dry matter | 91.2 ± 0.1 | 91.2 ± 0.5 | 91.2 ± 0.4 |
| Crude protein | 15.8 ± 0.1 | 22.7 ± 0.3 | 29.6 ± 0.8 |
| Crude ash | 5.49 ± 0.2 | 6.32 ± 0.2 | 7.14 ± 0.2 |
| Crude fat | 6.65 ± 0.1 | 7.09 ± 0.4 | 7.62 ± 0.4 |
| Neutral detergent fiber | 8.73 ± 0.1 | 11.4 ± 1.2 | 13.7 ± 1.9 |
| Acid detergent fiber | 2.40 ± 0.1 | 3.72 ± 0.6 | 4.02 ± 0.4 |
| Acid detergent lignin | 0.58 ± 0.1 | 0.99 ± 0.4 | 0.68 ± 0.3 |
| Titanium dioxide | 0.54 ± 0.1 | 0.56 ± 0.1 | 0.55 ± 0.1 |
| Alanine | 0.69 ± 0.1 | 0.97 ± 0.01 | 1.25 ± 0.03 |
| Arginine | 1.25 ± 0.1 | 1.77 ± 0.04 | 2.28 ± 0.07 |
| Aspartic Acid | 1.28 ± 0.1 | 2.06 ± 0.04 | 2.79 ± 0.08 |
| Cystine | 0.20 ± 0.1 | 0.30 ± 0.01 | 0.39 ± 0.01 |
| Glutamic Acid | 2.00 ± 0.1 | 3.21 ± 0.06 | 4.39 ± 0.11 |
| Glycine | 0.54 ± 0.1 | 0.82 ± 0.01 | 1.10 ± 0.03 |
| Histidine | 0.33 ± 0.1 | 0.50 ± 0.01 | 0.65 ± 0.02 |
| Isoleucine | 0.56 ± 0.1 | 0.85 ± 0.01 | 1.14 ± 0.03 |
| Leucine | 1.26 ± 0.1 | 1.76 ± 0.02 | 2.26 ± 0.06 |
| Lysine | 1.18 ± 0.1 | 1.59 ± 0.03 | 1.99 ± 0.05 |
| Methionine | 0.41 ± 0.1 | 0.49 ± 0.01 | 0.58 ± 0.01 |
| Methionine + Cysteine | 0.61 ± 0.1 | 0.79 ± 0.02 | 0.97 ± 0.02 |
| Phenylalanine | 0.68 ± 0.1 | 1.03 ± 0.02 | 1.35 ± 0.04 |
| Proline | 0.82 ± 0.1 | 1.15 ± 0.02 | 1.49 ± 0.04 |
| Serine | 0.64 ± 0.1 | 0.98 ± 0.01 | 1.31 ± 0.03 |
| Threonine | 0.77 ± 0.1 | 1.02 ± 0.01 | 1.27 ± 0.03 |
| Tryptophan | 0.18 ± 0.1 | 0.26 ± 0.01 | 0.35 ± 0.01 |
| Valine | 0.67 ± 0.1 | 0.98 ± 0.02 | 1.29 ± 0.04 |

### 2.4 statistical analyses and calculations

Apparent prececal digestibility coefficients (DC) for CP, sum of essential amino acids (SEAA), sum of non-essential amino acids (SNEAA), as well as for individual AA were calculated using the formula:

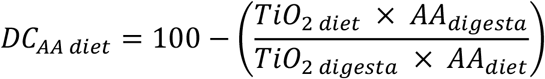

TiO_2 diet_ and TiO_2 digesta_ represent the dry matter concentrations of TiO_2_ in the diet and digesta and AA_diet_ and AA_digesta_ are the dry matter concentration of AA in diet and digesta.

The obtained DCs were then used to estimate the apparent prececal CP and AA digestibilty in regard to the dietary ingested amount of AA at given inclusion level of respective ESBM variants. The partial prececal AA digestibility from each ESBM was estimated by linear regression technique, using the slope of apparent ileal AA flux at the terminal ileum in relation to the respective increase in AA intake in response to rising dietary contents of respective ESBM variants (Rodehutscord et al., 2004). Additionally, the ratio of sulfuric AA to lysine was calculated to indicate changes in the biological value and quality of the true protein from different ESBM variants.

For statistical analyses, calculated mean values over single pens within experimental runs yielded to a total of 546 data points. For each experimental diet, a mean value was calculated including the respective six replicate values. In this way, 91 values for zootechnical parameters in response to different diets and each 45 data points for the apparent prececal CP and AA digestibilty of each ESBM variant were calculated. These values were used for linear regression analysis (y = a + bx) applying the variables TIA, KOH-soluble CP and reactive lysine, respectively (The R Project, Version 3.6.1). The threshold of significance was as assumed at p ≤ 0.05. Finally, each of 546 pen-wise mean values of partial digestibility data was further used for descriptive statistics.

Our experimental design was planned to reach in any case a minimum statistical power of 1-β=0.8. The respective power analysis was performed with G*Power 3.1.9.7 ( Faul et al., 2007; Faul et al., 2009) applying the dataset of Hoffmann et al. (2019) for the determination of the effect size at assumed α = 0.05.

## 3. Results

### 3.1 Zootechnical performance

Broilers were healthy throughout the experimental phase and mortality was <1% and did not correlate to dietary treatments.

In total, zootechnical performance varied considerably between individual pens. Live weight (LW) at the end of the experimental phase (d22) varied from 752g to 985g, Total weight gain (TWG) and feed conversion ratio (FCR) ranged from 313g and 1.50 in the group with the highest dietary TIA (8.5 mg/g) to 539g and 1.19 in the group with the lowest dietary TIA (0.4 mg/g).

As shown in figure 1 and table 5, rising dietary TIA negatively affected final LW, TWG, total feed intake (TFI) as well as FCR in a straight linear manner (p < 0.001). Accordingly, a stepwise increase of dietary TIA of 1 mg/g depressed LW, TWG, and TFI by 15.0 g, 16.5 g and 5.7 g, and increased FCR by 0.015.

**Figure 1:**
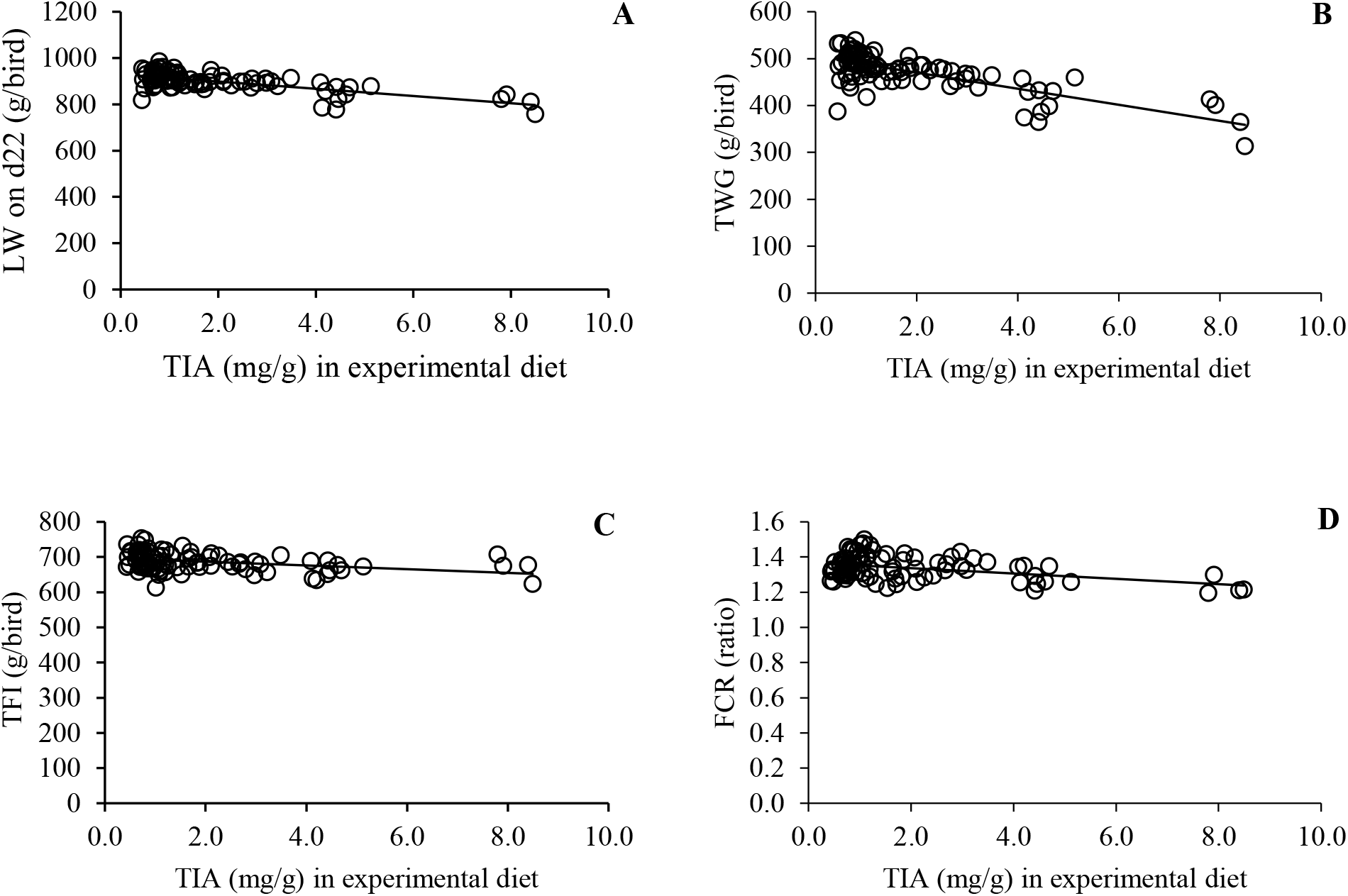
Effect of trypsin inhibitor activity (TIA) in mg/g in experimental diets on Live weight (LW) on d22 in g/bird (A), Total weight gain (TWG) in g/bird (B), total feed intake (TFI) in g/bird (C) and feed conversion ratio (FCR) as the ratio of consumed feed and live weight (D).

**Table 5:** Effect of trypsin inhibitor activity (mg/g) in experimental diets on live weight on d22 (g/bird), total weight gain (g/bird), total feed intake (g/bird) and feed conversion ratio (feed: gain) using linear regression models (y = a + bx; x = TIA)

|  | Regression model | Estimates of a | Estimates of b | R<sup>2</sup> |
| --- | --- | --- | --- | --- |
| Live weight (LW) on d22 g/bird | $y = a + bx$ | a, $930 \pm 4.4$ ($p < 0.001$) | b, $-15.0 \pm 1.8$ ($p < 0.001$) | 0.45 |
| Total weight gain (TWG), g/bird | $y = a + bx$ | a, $504 \pm 4.2$ ($p < 0.001$) | b, $-16.5 \pm 1.7$ ($p < 0.001$) | 0.63 |
| Total feed intake (TFI), g/bird | $y = a + bx$ | a, $698 \pm 3.9$ ($p < 0.001$) | b, $-5.7 \pm 1.6$ ($p < 0.001$) | 0.13 |
| Feed conversion ratio (FCR) | $y = a + bx$ | a, $1.37 \pm 0.009$ ($p < 0.001$) | b, $0.015 \pm 0.004$ ($p < 0.001$) | 0.17 |

The dietary amounts of KOH-soluble CP as well as the reactive lysine had no significant effect on zootechnical performance whatsoever (data not shown).

### 3.2 Partial prececal amino acid digestibility from different ESBM variants

Table 6 presents descriptive statistics of prececal digestibility of AA and CP arising from the ingestion of different ESBM variants. Like zootechnical performance, partial prececal digestibility varied noticeably with rising TIA levels. DC of CP, SEAA and SNEAA varied from 30.33% to 97.21%, 26.76% to 95.12% and 31.63% to 93.25%, respectively.

**Table 6:** Descriptive statistics of prececal digestibility of expeller extracted soybean meal variants (%) of amino acids, crude protein, sum of essential amino acids and sum of non-essential amino acids

|  | Mean | Standard Deviation | Median | 25% Quantil | 75% Quantil | Minimum | Maximum |
| --- | --- | --- | --- | --- | --- | --- | --- |
| Crude Protein | 74.29 | 14.34 | 79.50 | 64.10 | 86.10 | 30.33 | 97.21 |
| Sum of essential amino acids | 75.60 | 14.67 | 81.28 | 65.95 | 86.56 | 26.76 | 95.12 |
| Sum of non-essential amino acids | 73.53 | 13.38 | 77.84 | 64.35 | 83.78 | 31.63 | 93.25 |
| Alanine | 72.30 | 16.56 | 78.73 | 61.77 | 84.21 | 21.64 | 94.21 |
| Arginine | 82.64 | 10.33 | 86.66 | 76.37 | 90.02 | 44.84 | 96.51 |
| Aspartic Acid | 72.27 | 12.72 | 75.27 | 63.95 | 82.38 | 34.87 | 91.45 |
| Cystine | 50.49 | 19.23 | 54.41 | 36.22 | 67.14 | 4.94 | 83.49 |
| Glutamic Acid | 79.10 | 11.60 | 83.47 | 71.45 | 87.68 | 39.58 | 95.89 |
| Glycine | 68.40 | 14.50 | 71.86 | 58.70 | 80.18 | 25.18 | 90.73 |
| Histidine | 74.97 | 13.17 | 79.73 | 67.36 | 84.68 | 29.63 | 92.64 |
| Isoleucine | 73.71 | 16.95 | 80.18 | 64.10 | 85.89 | 17.68 | 95.51 |
| Leucine | 73.72 | 18.49 | 80.99 | 65.17 | 86.44 | 15.15 | 96.11 |
| Lysine | 77.34 | 12.83 | 81.36 | 70.26 | 87.59 | 34.66 | 94.71 |
| Methionine | 74.75 | 17.54 | 81.82 | 64.66 | 87.22 | 17.76 | 94.55 |
| Methionine + Cystine | 62.00 | 18.17 | 67.69 | 48.97 | 76.43 | 10.92 | 89.18 |
| Phenylalanine | 77.52 | 15.36 | 84.57 | 68.12 | 87.75 | 27.75 | 98.34 |
| Proline | 69.69 | 14.96 | 74.98 | 59.87 | 80.96 | 23.99 | 91.50 |
| Serine | 71.95 | 15.41 | 77.53 | 62.71 | 83.41 | 26.71 | 93.46 |
| Threonine | 67.97 | 15.08 | 72.49 | 57.64 | 80.58 | 23.93 | 90.62 |
| Tryptophan | 69.86 | 14.96 | 74.47 | 60.28 | 81.37 | 25.07 | 90.42 |
| Valine | 72.57 | 16.49 | 78.90 | 64.20 | 84.55 | 18.03 | 94.09 |

According to figure 2 and table 7., prececal digestibility of all individual AA as well as of CP of ESBM variants was significantly affected by the respective TIA level in a straight linear fashion (p < 0.001). On average, each increase of TIA by 1 mg/g reduced digestibility of CP as well as sums of essential AA and non-essential AA by 1.84%, 1.99% and 1.75%, respectively. The magnitude of TIA effects on AA digestibility differed between individual AA. Arginine digestion was least affected by TIA with an average value for all used ESBM of 82% but ranged from 44.8% to 96.5%. In contrast, the digestibility of cystine was impaired the most by increasing dietary TIA levels, with only 10.59% of prececal digestibility when feeding raw ESBM (23.6 mg/g TIA). The measured maximum of prececal cystine digestibility was at 83% (0.3 mg/g TIA), which also fell below the maxima of all other AA (≥ 89%). Linear regression models were established to quantify the impact of TIA on individual AA digestibility (table 7), which decreased in any case significantly (p < 0.001) with increasing levels of TIA. In figure 2., the effect of TIA on digestibility of CP, SEAA, SNEAA, lysine, methionine and cystine is illustrated. Furthermore, the ratio of digestible sulfur containing AA to lysine was calculated to characterize the impact of TIA on the biological value of the digestible AA (figure 3). Per unit increase of TIA in ESBM, the ratio of ileal digested sulfuric AA to lysine significantly decreased by 0.0002 (from ~0.9 down to ~0.5) (p < 0.001).

**Figure 2:**
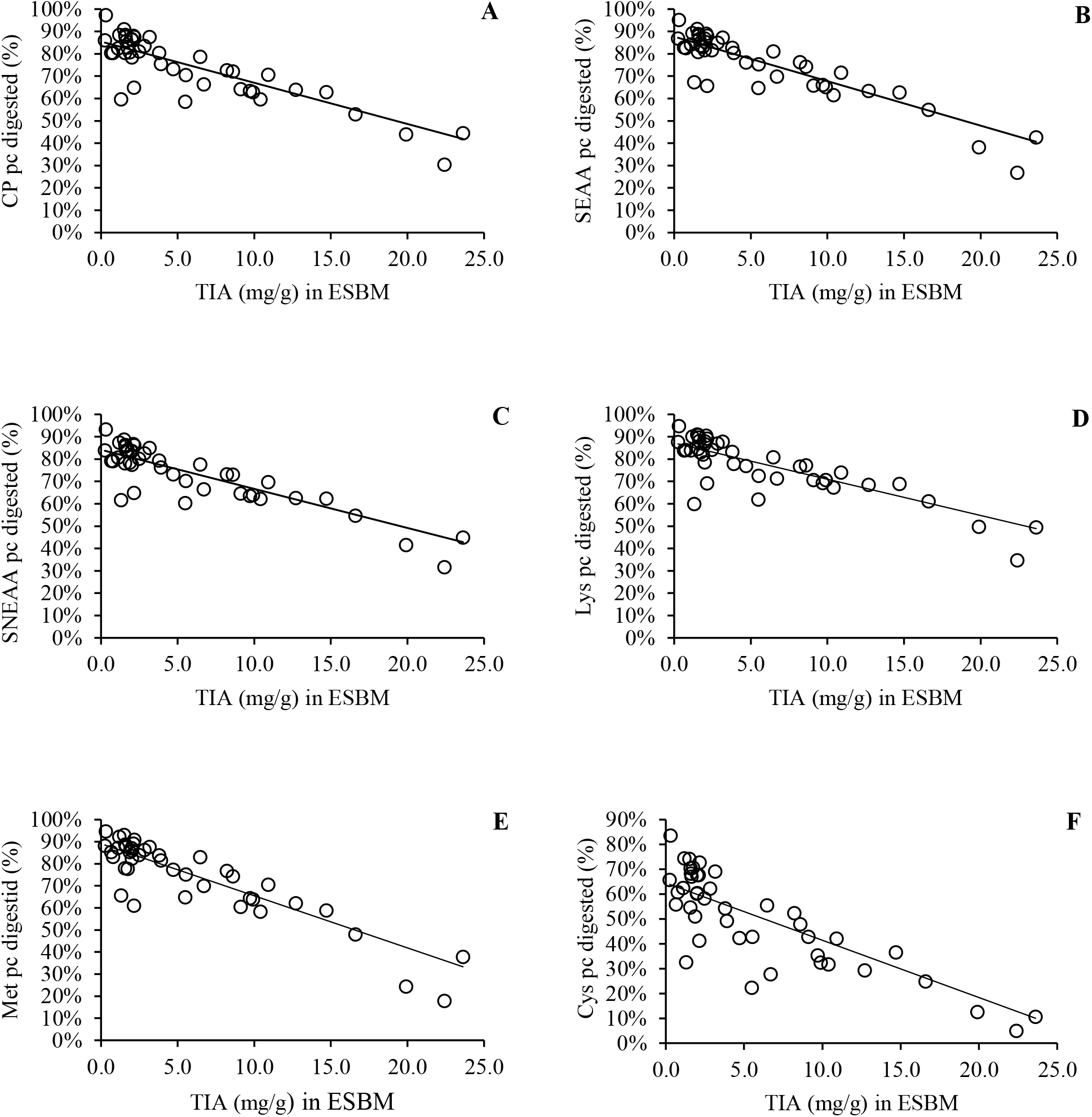
Effect of trypsin inhibitor activity (TIA) in mg/g on percentage of prececal (pc) digested crude protein (CP) (A), sum of essential amino acids (SEAA) (B), sum of non-essential amino acids (SNEAA) (C), lysine (Lys) (D), methionine (Met) (E) and cystine (Cys) (F) of expeller extracted soybean meal (ESBM).

**Table 7:** Effect of trypsin inhibitor activity (mg/g) in expeller extracted soybean meals on partial prececal digestibility of crude protein, sum of essential amino acids, sum of non-essential amino acids and individual amino acids using linear regression models (regression model: y = a + bx; x = TIA)

|  | Regression models | Estimates of a | Estimates of b | R<sup>2</sup> |
| --- | --- | --- | --- | --- |
| Crude Protein | $y = a + bx$ | a, $85.48 \pm 1.85$ ($p < 0.001$) | b, $-1.84 \pm 0.21$ ($p < 0.001$) | 0.64 |
| Sum of essential amino acids | $y = a + bx$ | a, $87.68 \pm 1.70$ ($p < 0.001$) | b, $-1.99 \pm 0.20$ ($p < 0.001$) | 0.71 |
| Sum of non-essential amino acids | $y = a + bx$ | a, $84.14 \pm 1.68$ ($p < 0.001$) | b, $-1.75 \pm 0.19$ ($p < 0.001$) | 0.66 |
| Alanine | $y = a + bx$ | a, $85.75 \pm 1.98$ ($p < 0.001$) | b, $-2.22 \pm 0.23$ ($p < 0.002$) | 0.69 |
| Arginine | $y = a + bx$ | a, $91.10 \pm 1.21$ ($p < 0.001$) | b, $-1.39 \pm 0.14$ ($p < 0.001$) | 0.70 |
| Aspartic Acid | $y = a + bx$ | a, $81.91 \pm 1.73$ ($p < 0.001$) | b, $-1.59 \pm 0.20$ ($p < 0.001$) | 0.60 |
| Cystine | $y = a + bx$ | a, $64.49 \pm 2.76$ ($p < 0.001$) | b, $-2.31 \pm 0.32$ ($p < 0.001$) | 0.55 |
| Glutamic Acid | $y = a + bx$ | a, $88.32 \pm 1.45$ ($p < 0.001$) | b, $-1.52 \pm 0.17$ ($p < 0.001$) | 0.66 |
| Glycine | $y = a + bx$ | a, $79.62 \pm 1.91$ ($p < 0.001$) | b, $-1.85 \pm 0.22$ ($p < 0.001$) | 0.62 |
| Histidine | $y = a + bx$ | a, $85.58 \pm 1.61$ ($p < 0.001$) | b, $-1.75 \pm 0.19$ ($p < 0.001$) | 0.68 |
| Isoleucine | $y = a + bx$ | a, $87.91 \pm 1.88$ ($p < 0.001$) | b, $-2.34 \pm 0.22$ ($p < 0.001$) | 0.73 |
| Leucine | $y = a + bx$ | a, $89.27 \pm 2.03$ ($p < 0.001$) | b, $-2.56 \pm 0.24$ ($p < 0.001$) | 0.74 |
| Lysine | $y = a + bx$ | a, $87.18 \pm 1.71$ ($p < 0.001$) | b, $-1.62 \pm 0.20$ ($p < 0.001$) | 0.61 |
| Methionine | $y = a + bx$ | a, $89.08 \pm 2.07$ ($p < 0.001$) | b, $-2.36 \pm 0.24$ ($p < 0.001$) | 0.70 |
| Methionine + Cystine | $y = a + bx$ | a, $76.31 \pm 2.31$ ($p < 0.001$) | b, $-2.36 \pm 0.27$ ($p < 0.001$) | 0.65 |
| Phenylalanine | $y = a + bx$ | a, $90.40 \pm 1.69$ ($p < 0.001$) | b, $-2.12 \pm 0.20$ ($p < 0.001$) | 0.74 |
| Proline | $y = a + bx$ | a, $81.63 \pm 1.86$ ($p < 0.001$) | b, $-1.97 \pm 0.22$ ($p < 0.001$) | 0.66 |
| Serine | $y = a + bx$ | a, $84.60 \pm 1.80$ ($p < 0.001$) | b, $-2.08 \pm 0.21$ ($p < 0.001$) | 0.70 |
| Threonine | $y = a + bx$ | a, $79.85 \pm 1.92$ ($p < 0.001$) | b, $-1.96 \pm 0.22$ ($p < 0.001$) | 0.65 |
| Tryptophan | $y = a + bx$ | a, $81.52 \pm 1.84$ ($p < 0.001$) | b, $-1.92 \pm 0.21$ ($p < 0.001$) | 0.66 |
| Valine | $y = a + bx$ | a, $86.11 \pm 1.93$ ($p < 0.001$) | b, $-2.23 \pm 0.22$ ($p < 0.001$) | 0.70 |

**Figure 3:**
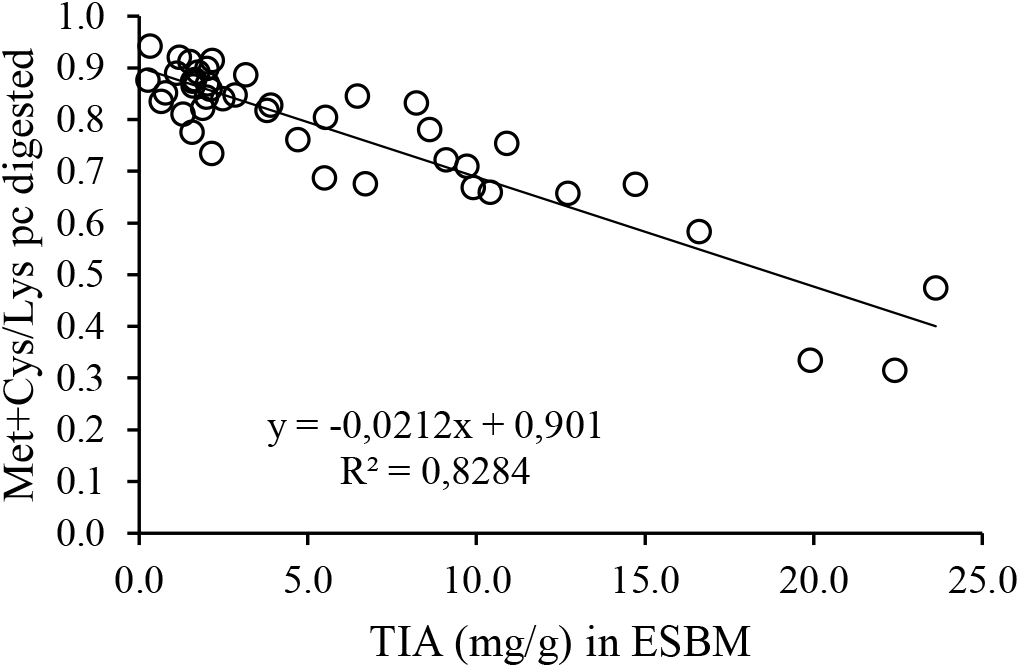
Effect of trypsin inhibitor activity (TIA) in mg/g on the ratio of prececal (pc) digested methionine + cystine (Met + Cys) and lysine in expeller extracted soybean meal (ESBM).

Prececal digestibility of CP and AAs was not significantly affected by the amount of KOH-soluble CP or reactive lysine, respectively (data not shown).

## 4. Discussion

In recent years, there have been several studies concerning the negative effect of TIA on growth performance of different animal models including rats (Grant et al., 1995), pigs (Batterham et al., 1993; Herkelman et al., 1992; Zollitsch et al., 1993), chickens (Clarke and Wiseman, 2007, 2005) and turkeys (Mian and Garlich, 1995). All these studies gained comparable results: the lower TIA in feed, the better growth performance. Consequently, soybeans are treated with heat and pressure to reduce this anti-nutritional potential to ensure proper performance and animal wellbeing. Clarke and Wiseman (2007, 2005) claim in their studies to reduce the TIA in full-fat soybeans to 4.0 mg/g is sufficient for broilers. Nevertheless, an overtreatment of soybeans can also lead to decreased growth performance (Araba and Dale, 1990; Pacheco et al., 2014). In the present study, the solubility of CP in KOH varied from 64.4% to 97.7% of total CP and reactive lysine ranged from 14.7% to 25.0% of DM. Neither CP solubility in KOH, nor reactive lysine correlated with growth performance or AA digestibility. This is in good agreement to earlier published data of Herkelman et al. (1991), who gained in their trials the highest chick performance at a minimum of 50% protein solubility. Furthermore, Hoffmann et al. (2019) fed experimental diets comparable to ours during a whole fattening trial with broiler chickens and did not observe any negative effects of KOH-soluble CP or reactive lysine whatsoever. Therefore, we conclude that protein denaturation by heat treatment of ESBM variants from the present study was negligible and TIA was the dominant antinutritional factor modulating zootechnical performance and protein digestibility.

The aforementioned study of Hoffmann et al. (2019), observed a straight linear decrease of zootechnical performance and especially feeding efficiency of broiler chicks when fed diets with finely graded differences in dietary TIA ranging from 0.3-8.7 mg/g. In fact, using ESBM with TIA below 4.0 mg/g further improved feed efficiency of up to 16% until the end of the grower stage. In the present study, we observed a significant linear reduction in LW and TWG. Consistent with Hoffmann et al. (2019), this effect responded in a straight linear fashion indicating further improvement of production efficiency when decreasing TIA in full-fat soybeans below 4.0 mg/g. However, TFI and FCR were less impaired by TIA than LW and TWG, which seems to contradict earlier data at first glance. However, the experimental phase in our trials was set from day 15 to day 22 according to the experimental model of Rodehutscord et al. (2004) whereas Hoffmann et al. (2019) observed the zootechnical performance throughout the whole grower and finisher stages of fattening. This may explain why TFI and FCR was less responsive in the present study. Clarke and Wiseman (2007), who recorded weight gain and feed intake for only 3 days observed also decreased weight gain with rising TIA levels but no significant correlation to feed intake. They concluded, that the recording phase of these parameters have to be at least 21 days to gain stable results.

For the estimation of feed protein quality, the determination of prececal digestibility is the most common method today. The method is based on the idea of measuring the “unabsorbed” AA directly in the terminal ileum and estimating the product specific digestibility by calculating the slope of increasing apparently digested amounts of AA at the terminal ileum over a graded increase in the intake of product-specific AA. The advantage of this method is that digesta are collected directly from the terminal ileum, which means there is negligible bias by microbial fermentation and no contamination with renal excretions and other materials. In addition, since the slope predominantly reflects the product-specific AA digestibility, there is no need to correct the data for the endogenously secreted amounts of AA. This has been demonstrated by Rodehutscord et al. (2004), who observed a linear relationship between product-specific AA intake and quantitative AA flow at the terminal ileum. Kluth et al. defined in 2005 the section of the intestine which needs to be collected. Regarding recent literature, many authors have used this method for estimating protein quality in different feedstuff for broiler chickens (Short et al., 1999, Kluth and Rodehutscord, 2009, Foltyn et al., 2015, Rada et al., 2017).

In the present study, AA digestibility from individual ESBM products showed the same linear response to TIA as growth performance. The higher TIA in the respective soybean product, the lower the associated digestibility. TIA affected the apparent digestibility of every individual AA. The DC of arginine, glutamic acid, phenylalanine and lysine were the least affected AA. In contrary to those, cystine and methionine showed a markedly increased responsiveness to TIA in ESBM. At 0.3 mg/g TIA 83.5% of cystine were apparently digested in the terminal ileum, whereas at TIA of 22.6 mg/g only 4.9% of cystine was absorbed. This is in good agreement with the findings of Clarke and Wiseman (2007). They found out that DC of cysteine and methionine showed the strongest correlation to TIA. In their trials, DC varied from 71.8% at 1.9 mg/g to 34.5% at 14.8 mg/g.

One explanation for the negative impact of elevated dietary TIA levels on AA digestibility could be that TI bind irreversibly on the digestive enzymes trypsin and chymotrypsin and thereby impair protein digestion. Foltyn et al. (2015) measured the trypsin activity in the jejunum and discovered a reduction in enzyme activity when feeding raw full fat soybeans to chickens for 4 days. This period is comparable to that from the present study. Hence, it appears plausible that the activity of trypsin and chymotrypsin were negatively affected by KTI and BBI from ESBM of the present study. This conclusion may vary under experimental conditions comprising longer periods of treatment feeding, since the organism tends to adapt over time to the antinutritive stimulus by increased pancreatic secretion to satiate the binding sites of the inhibitor pool and provide a surplus of active trypsin and chymotrypsin (Lyman and Lepkovski, 1957; Nitsan and Liener, 1976). In addition, BBI from soybeans contain many disulfide bonds and are consequently rich in cysteine (Odani and Ikenaka, 1973). Accordingly, the higher amounts of sulfur-containing AA at the terminal ileum may also derive from an enrichment of BBI with further increase in the dietary proportion of ESBM with high TIA.

Besides that, high endogenous losses of cystein and methionine may further explain the high concentrations of sulfuric AA at the terminal ileum. Trypsin and chymotrypsin are rich in sulfur-containing AA (Nitsan and Liener, 1976). Peptide hormone CCK promotes the exocrine pancreatic synthesis and secretion of the digestive enzymes in response to increasing TIA in the small intestinal lumen. Fölsch et al. (1978) reported that continuous injections with CCK increased the pancreatic weight of rats. They could also prove a hypertrophy and hyperplasia of the gland. Furuse et al. (1990) observed in their survey a prompt increase of plasma CCK after 15 minutes, when feeding rats with soybean trypsin inhibitor. In the present study, we used the same ESBM as Hoffmann et al. (2019). In their feeding trials, a linear increase of pancreatic weight with rising TIA levels in feed was observed at the end of the finishing phase. Therefore, it is plausible that there was also increased activity of the exocrine pancreas in the present study, although presumably to a lower extend given the short experimental duration. At this point, it is not possible to estimate, which of the above discussed potential causes of the reduced digestibility of sulfuric AA had the strongest effect on apparent prececal AA digestibility. Hence, this issue must be addressed in follow-up studies.

Increasing TIA from ESBM did not just decrease prececal protein utilization from a quantitative perspective but also affected the qualitative value of the available protein. Specifically, the ratio of apparently digested sulfuric AA to lysine decreased linearly with increasing TIA from ~0.9 to ~0.5, which was due to the aforementioned disproportionately higher digestive depression for cystine compared to other AA and especially lysine. In other terms, the degree of TIA reduced the biological value of the absorbed true protein. The biological value of feed protein reflects how close the spectrum of amino acids represents the ideal protein for growing organisms. The closer the amino acid spectrum in feed resembles the ideal protein at a given stage of the production cycle, the lesser total CP and true protein is necessary to meet the animals demand (van Milgen and Dourmad, 2015). Hence, an increasing dietary TIA by usage of poorly processed ESBM further increases the necessity to either increase total ESBM in complete feed or to balance the AA spectrum by adding crystalline AA. Both strategies increase feed costs and thus impair the economic success of poultry production.

In conclusion, varying dietary TIA from the usage of differentially processed ESBM variants negatively affected zootechnical performance and prececal AA digestibility in a straight linear fashion. Every single AA was affected but cysteine showed the lowest values over the whole range of applied TIA. To the best of our knowledge, this is the first study to show significant effects of TIA <4.0 mg/g in soy products on prececal AA digestion. The linearity of the observed effects questions the suitability of defined upper limits for TIA in soybean products. In contrast, the KOH-soluble CP as well as the amounts of reactive lysine in respective diets did not affect the investigated parameters. This suggests, that TIA could be decreased close to zero under the present conditions without promoting negative effects through excessive denaturation or formation of maillard products. In addition to the quantitative effects of TIA on feed protein utilization, it was further observed that the quality of the digestible spectrum of AA decreased significantly. This was evident by a marked change in the ratio of sulfuric AA to lysine. Overall, these findings suggest that the TIA of dietary components should be considered when determining their protein value *in vivo*.

## Abbreviations

AA: amino acid
BBI: Bowman-Birk trypsin inhibitors
CCK: Cholecystokinin
CP: crude protein
DC: apparent prececal digestibility coefficient
ESBM: expeller extracted soybean meal
FCR: feed conversion ratio
KOH: potassium hydroxide
KTI: Kunitz trypsin inhibitors
LW: live weight
SEAA: sum of essential amino acids
SNEAA: sum of non-essential amino acids
TFI: total feed intake
TI: trypsin inhibitor
TIA: trypsin inhibitor activity
TiO_2_: titanium dioxide
TWG: total weight gain

## Author contributions

**S. Kuenz:** Formal analysis, Investigation, Writing – Original Draft, Visualization, Project administration **S. Thurner:** Conceptualization, Funding acquisition, Resources, Project administration, Supervision **D. Hoffmann:** Conceptualization, Investigation **K. Kraft:** Conceptualization, Investigation **M. Wiltafsky:** Investigation **K. Damme:** Resources **W. Windisch:** Conceptualization, Writing – Review & Editing, Supervision **D. Brugger:** Conceptualization, Writing – Review & Editing, Supervision.

## Acknowledgements

The project was supported by funds of the Federal Ministry of Food and Agriculture (BMEL) based on a decision of the Parliament of the Federal Republic of Germany via the Federal Office for Agriculture and Food (BLE) under the innovation support program (Grant number: 2814EPS022). Thanks to the staff of the Chair of Animal Nutrition, the Department for Education and Poultry Research and the Institute of Agricultural Engineering and Animal Husbandry for excellent technical assistance. The authors are especially grateful to Prof. Rodehutscord (Hohenheim University) and his team for the mixing of the experimental diets.

